# Transposable Elements Facilitate the De Novo Origin of Antifreeze Protein and the Diversification of Its Gene Family in Snailfishes

**DOI:** 10.64898/2026.04.28.721326

**Authors:** Nathan Rives, Prabodh Bajpai, Xuan Zhuang

## Abstract

Transposable elements (TEs) are increasingly recognized as important sources of genomic innovation, yet mechanistically resolved examples of how they help generate new functional genes in vertebrates remain rare. Type I antifreeze proteins (AFPI) in fishes are life-saving adaptations shaped by strong freezing selection and provide an exceptional system for studying new gene evolution under extreme environmental pressure. We recently showed that AFPI in flounder, cunner, and sculpin evolved independently through distinct partial de novo routes, converging on a nearly identical alanine-rich antifreeze protein. Here, we elucidate the origin and evolution of AFPI in the last remaining unresolved lineage, snailfishes, using a chromosome-scale genome assembly for *Liparis atlanticus* together with multi-tissue Iso-Seq, tissue-specific RNA-seq, and comparative genomics across AFPI-bearing and AFPI-lacking snailfishes and teleost outgroups. We show that snailfish AFPI originated within *Liparis* and rapidly diversified as a young gene family with multiple isoforms and lineage- and population-specific copy-number change. Genome-wide homology searches support a de novo origin of the alanine-rich coding region from noncoding sequence rather than from a pre-existing protein-coding precursor. In contrast, the surrounding regulatory architecture was assembled through sequence recruitment: a hAT-derived fragment contributes promoter- and transcription-start-site-proximal sequence, and a conserved noncoding segment together with a Ty3/Gypsy-derived long terminal repeat (LTR) contributes the 3′ regulatory region. TE-rich locus structure also provides plausible mechanisms for subsequent locus expansion and translocation. Together, these results reveal a TE-facilitated, mosaic route to new gene evolution in vertebrates, demonstrating how noncoding DNA, repetitive sequence, and TE-derived regulatory fragments can be assembled into a strongly selected adaptive innovation.

**Author Summary:** Where do new genes with brand-new functions come from? We tackled this question using one of evolution’s clearest natural experiments: antifreeze proteins, life-saving molecules favored by selection because fish without them freeze in icy seawater. In this study, we show that mobile DNA called transposable elements helped build a new antifreeze gene in stages. Different transposable elements appear to have played different roles: one helped switch on a previously silent stretch of noncoding DNA, others contributed control sequences at the beginning and end of the gene, and repeat-rich DNA around the locus likely promoted gene duplication, movement to a new chromosomal location, and rapid diversification into a gene family. This is an unusually clear vertebrate example of how a new gene can emerge not in a single leap, but through stepwise assembly from different pieces of the genome. More broadly, our work shows that transposable elements do much more than disrupt genomes. Under strong natural selection, they can help turn noncoding DNA into a life-saving adaptation and then help that innovation expand and diversify.

## Introduction

Novel genes are fundamental drivers of biological innovation, yet the origins and mechanisms by which they arise are often only partially resolved. Beyond classical gene duplication [1], more than a dozen distinct molecular mechanisms for generating new genes have now been identified, including exon shuffling, retroposition, gene fusion and fission, horizontal gene transfer, de novo origination, or transposable element (TE) domestication [2]. The discovery of these diverse routes has greatly expanded our view of how genomes generate new functions and has revealed that mechanisms once thought improbable can, in fact, be repeatedly exploited across the tree of life. Many recently identified genes lack obvious protein coding precursors, indicating that additional, sometimes idiosyncratic evolutionary trajectories remain to be uncovered.

Among these mechanisms, the de novo birth of a gene from a previously noncoding sequence, while once thought to be highly unlikely, has now been clearly demonstrated across diverse taxa [3–7]. De novo genes are typically defined as genes that originate from ancestrally non-genic sequence that has newly acquired both transcriptional activity and an open reading frame (ORF) [4]. Such novel coding regions can arise from diverse substrates, including intergenic sequence, existing expressed enhancers, and long noncoding RNAs, or by gain of function in existing noncoding genes [7]. Because these substrates and routes to transcriptional activation and ORF formation are so diverse, it is often possible to recognize that a gene is de novo in origin while still lacking a mechanistic reconstruction of how its coding and regulatory components were assembled. This gap motivates detailed studies that trace how particular de novo genes recruit, combine, and modify genomic elements to form a functional gene.

TEs are now recognized as important contributors to genetic novelty and new gene formation, providing raw material for both coding and regulatory innovation [8–11]. At the coding level, TE-derived sequences can be domesticated and co-opted for a host trait [12], exapted and modified to confer new functions [11, 13] or mediate RNA-based retroposition of transcribed genes, producing retrogenes with potentially new function [14–16]. At the regulatory level, TE insertions in intronic or intergenic regions can supply cis-regulatory elements that modulate gene expression [17]. Beyond modifying existing genes, recent work further shows that de novo expressed ORFs (neORFs), precursors of de novo genes, frequently arise in TE-associated genomic regions [18], and that TE insertions can promote the emergence of new transcripts by introducing transcription factor–binding motifs [19]. Together, these observations highlight TE-rich regions as fertile ground for the coding and regulatory components of new genes, yet we still lack mechanistically detailed examples of how TE-derived fragments are assembled into a specific, functional de novo gene.

Fish antifreeze proteins (AFPs) provide a natural model for studying how new genes originate and acquire essential functions under strong selection. AFPs evolved in response to marine glaciation and are essential for the survival of polar and subpolar fishes living in waters below the freezing point of their body fluids, where they would otherwise die of inoculative freezing [20–22]. By binding to nascent ice crystals and inhibiting their growth, AFPs prevent lethal internal ice formation [21]. Across multiple, phylogenetically distinct lineages, AFPs have independently evolved into four structurally unrelated types—AFPI, AFPII, AFPIII, and antifreeze glycoproteins (AFGP)—a striking example of functional convergence [22, 23]. Work on these four classes has revealed diverse evolutionary routes to new genes within a single adaptive system, including partial and fully de novo gene birth [24–26], neofunctionalization of a preexisting enzyme [27], and origin from an existing gene followed by horizontal gene transfer [28–30]. This combination of independent origins, well-defined life-saving function, and relatively recent evolutionary timing makes AFPs a powerful system for investigating mechanisms of new gene evolution.

Among these AFPs, type I antifreeze proteins (AFPI) are particularly interesting. AFPI occurs in four distantly related fish lineages: flounder, cunner, sculpin, and snailfish, spanning three orders and four families [31–34]. Despite this deep phylogenetic separation, AFPI proteins in these lineages share remarkably similar sequences and structures: small, amphipathic α helices composed of alanine-rich sequence with periodic threonine [35–37]. Our recent comparative genomic work showed that, rather than descending from a single ancestral AFPI gene, AFPI in flounder, cunner, and sculpin arose independently from distinct progenitor genes via partial de novo mechanisms, providing a rare example of convergent evolution at the protein sequence level [38]. AFPI therefore offers an opportunity to study how genes with the same function and highly similar sequences can emerge multiple times from different genomic substrates under similar selective pressures. Its relatively recent origin, likely during Miocene marine glaciation within the last 10 to 20 million years [39], further increases the likelihood that progenitor sequences remain detectable in extant genomes. Among the four AFPI-bearing lineages, snailfish AFPI is distinctive in that it has been linked to TE-rich genomic regions and shows little similarity to canonical protein-coding genes [40], yet its genomic precursors, locus organization, and evolutionary mechanisms have not been fully reconstructed.

Here, we investigate the genomic origins and evolutionary mechanisms of snailfish AFPI. We generated a high-quality long-read genome assembly for *Liparis atlanticus* and performed multi-tissue Iso-Seq and tissue-specific RNA-seq to characterize AFPI gene structure, isoforms, and tissue associated expression patterns. We then compared AFPI-bearing (AFPI+) and AFPI-lacking (AFPI–) snailfishes, as well as closely related outgroup genomes, to reconstruct AFPI-containing loci, identify conserved flanking genes, and infer the ancestral genomic components involved in AFPI formation, revealing two distinct loci with lineage-specific expansion and loss. Our genomic and transcriptomic analyses indicate that snailfish AFPI arose through a partial de novo process in which TE-derived and noncoding sequences were assembled into a functional antifreeze protein gene, with patterns of TE insertion and co-localization implicating TEs in locus seeding, regulatory control, copy number expansion, and relocation of AFPI copies.

Together, these results provide a mechanistically detailed example of TE facilitated de novo gene birth and subsequent diversification in a vertebrate and illustrate how multiple genomic sources can be co-opted to generate a new, strongly selected gene in extreme environments.

## Results

### Chromosome-level genome assembly and refinement of *Liparis atlanticus*

We generated a chromosome-level assembly of the Atlantic snailfish, *L. atlanticus,* using PacBio HiFi sequencing and Hi-C scaffolding. PacBio HiFi sequencing produced 3.14 million circular consensus reads totaling 60.2 Gb, corresponding to approximately 102× coverage of the final assembly. The initial assembly contained a high rate of duplicated BUSCO genes (∼64%), which may have been a result of the genome’s high heterozygosity (> 2%). Because Hifiasm did not fully collapse redundant haplotigs, we further refined the assembly with Purge_Dups, which substantially reduced contig number and lowered duplicated BUSCOs to 1.8%. The final assembly size was 585,533,586 bp, comparable to those of other sequenced snailfishes (Table 1). The *L. atlanticus* genome was then compared to two AFPI+ snailfish *Liparis gibbus* (GCA_040955725.1) and *Liparis tanakae* (GCA_050084785.1), one AFPI– snailfish *Pseudoliparis swirei* (GCF_029220125.1) [41], and a closely related AFPI– outgroup teleost fish *Cyclopterus lumpus* (GCF_009769545.1) [42] for downstream analyses in this study (Table 1, Supplementary Fig. 1).

**Table 1:** Genome assembly assessment.

| Species | *Liparis*<br>*atlanticus* | *Liparis*<br>*gibbus* | *Liparis*<br>*tanakae* | *Pseudoliparis*<br>*swirei* | *Cyclopterus*<br>*lumpus* |
| --- | --- | --- | --- | --- | --- |
| **Genome**<br>**assembly** |  |  |  |  |  |
| Scale | Chromosome | Contig | Chromosome | Chromosome | Chromosome |
| Total size | 586.1 Mb | 677.8 Mb | 574.4 Mb | 625.1 Mb | 572.9 Mb |
| Contigs | 1643 | 51279 | 1631 | 1102 | 395 |
| Scaffolds | 462 | NA | 125 | 199 | 48 |
| Chromosomes | 24 | NA | 24 | 24 | 25 |
| N50 | 25 Mb | 590.1 kb | 24.6 Mb | 25.7 Mb | 23.9 Mb |
| TE Content (%) | 32 | 34 | 30 | 36 | 21 |
| **BUSCO**<br>**Assessment** |  |  |  |  |  |
| Complete (%) | 95.6 | 96.1 | 95.7 | 96.7 | 98.1 |
| Complete and<br>single copy (%) | 93.8 | 95.0 | 92.9 | 95.2 | 97.3 |
| Complete and<br>duplicated (%) | 1.7 | 1.1 | 2.8 | 1.5 | 0.8 |
| Fragmented (%) | 1.0 | 1.9 | 1.6 | 1.0 | 0.5 |
| Missing (%) | 3.4 | 2.0 | 2.7 | 2.3 | 1.4 |

### Two syntenic AFPI loci and secondary-locus translocation

To identify AFPI gene repertoires and map their genomic loci in snailfishes, we used all AFPI transcripts recovered from Iso-Seq and RNA-seq data generated in this study, together with two GenBank AFPI mRNA sequences, as BLAST queries against the genomes of three AFPI+ species. Unlike the other three AFPI+ lineages, in each lineage all AFPI genes are confined to a single genomic locus [38], snailfishes harbor AFPI genes at two conserved syntenic genomic loci on different chromosomes: an *ETV6–PARP12* interval and a *TNS3–IGBF* interval (Fig. 1A). In *L. atlanticus*, we identified 62 AFPI genes, including six pseudogenes, all located between *ETV6* and *PARP12*. In *L. gibbus*, four AFPI genes occur in the *ETV6–PARP12* interval and two at *TNS3–IGBF* interval. In *L. tanakae*, three available assemblies (GCA_050084785.1; GCA_036178185.1; GCA_006348945.1)[43] from two different populations show marked variation, ranging from 0 to 2 AFPI genes per locus (Fig. 1B). Although AFPI gene number and locus occupancy vary among species and populations, the flanking genes at both loci are conserved in AFPI– outgroups.

**Figure 1.**
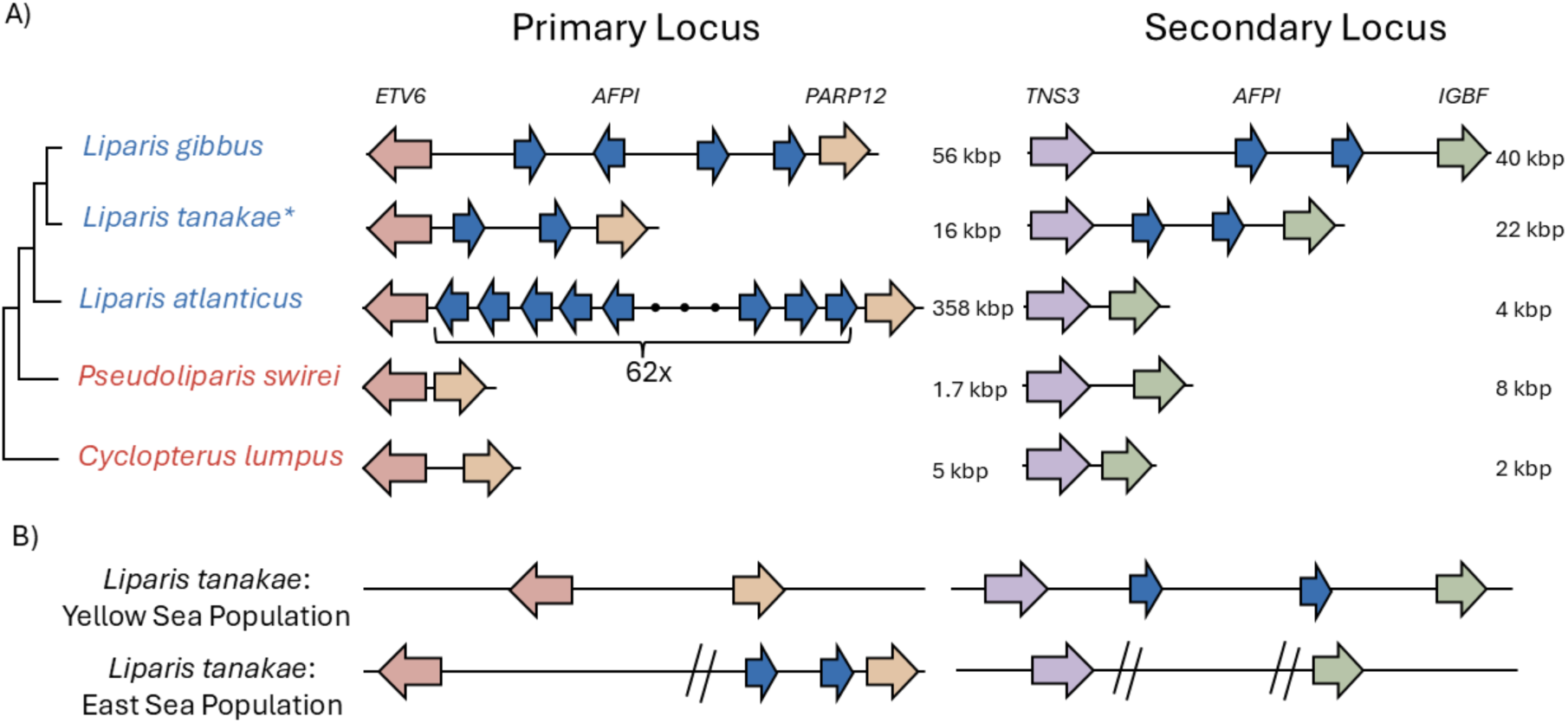
Genomic organization of AFPI loci in *Liparis* and outgroup genomes. (A) Organization of the primary locus, bounded by *ETVc* and *PARP12*, and the secondary locus, bounded by *TNS3* and *IGBF*, in three AFPI-bearing *Liparis* species, one AFPI-lacking snailfish, and one AFPI-lacking outgroup. Colored arrows indicate annotated genes and their orientation; arrow length is not drawn to scale. Locus sizes indicate the distance between the two flanking genes in each interval. (B) Comparison of representative *L. tanakae* assemblies from two populations, showing population-level variation in AFPI locus organization. Double slashes indicate assembly breaks. The East Sea assembly also contains five additional AFPI genes on unplaced contigs, which are not shown. The *L. tanakae* locus map in panel A combines the AFPI loci observed in the two population-specific assemblies shown in panel B. The species tree was inferred by maximum-likelihood analysis of a concatenated alignment of 2,940 single-copy BUSCO orthologs identified in the whole-genome assemblies of all five species.

We designate the *ETV6–PARP12* interval as the primary locus and the *TNS3–IGBF* interval as the secondary locus because the primary locus is the only AFPI locus in *L. atlanticus*, whereas AFPI at the secondary locus is absent from *L. atlanticus* but present in *L. gibbus* and some *L. tanakae* assemblies. This distribution indicates that the AFPI secondary locus arose after the divergence of *L. atlanticus*. AFPI genes at the secondary locus are also accompanied by duplicated sequence homologous to the 5′ upstream region and exon 1 of *PARP12*, features that characterize the primary AFPI locus but are not part of the native *TNS3–IGBF* interval. These shared flanking fragments support movement of an AFPI-containing genomic segment from the primary locus to the secondary locus, rather than independent origin of AFPI at the two sites.

### AFPI isoform diversity and tissue-associated expression

To classify AFPI isoforms, we compared amino acid sequences from the three AFPI+ snailfish species and used an AFPI gene tree as support for grouping related copies (Supplementary Fig. 2). Together, these data resolved three principal isoform classes, long, primary, and secondary, that differ in sequence length, nonrepetitive amino acid content, and genomic distribution (Fig. 2). The primary isoform is found only in the primary locus and comprises most AFPI copies. The long isoform occurs as a single copy in each species and is also restricted to the primary locus.

**Figure 2.**
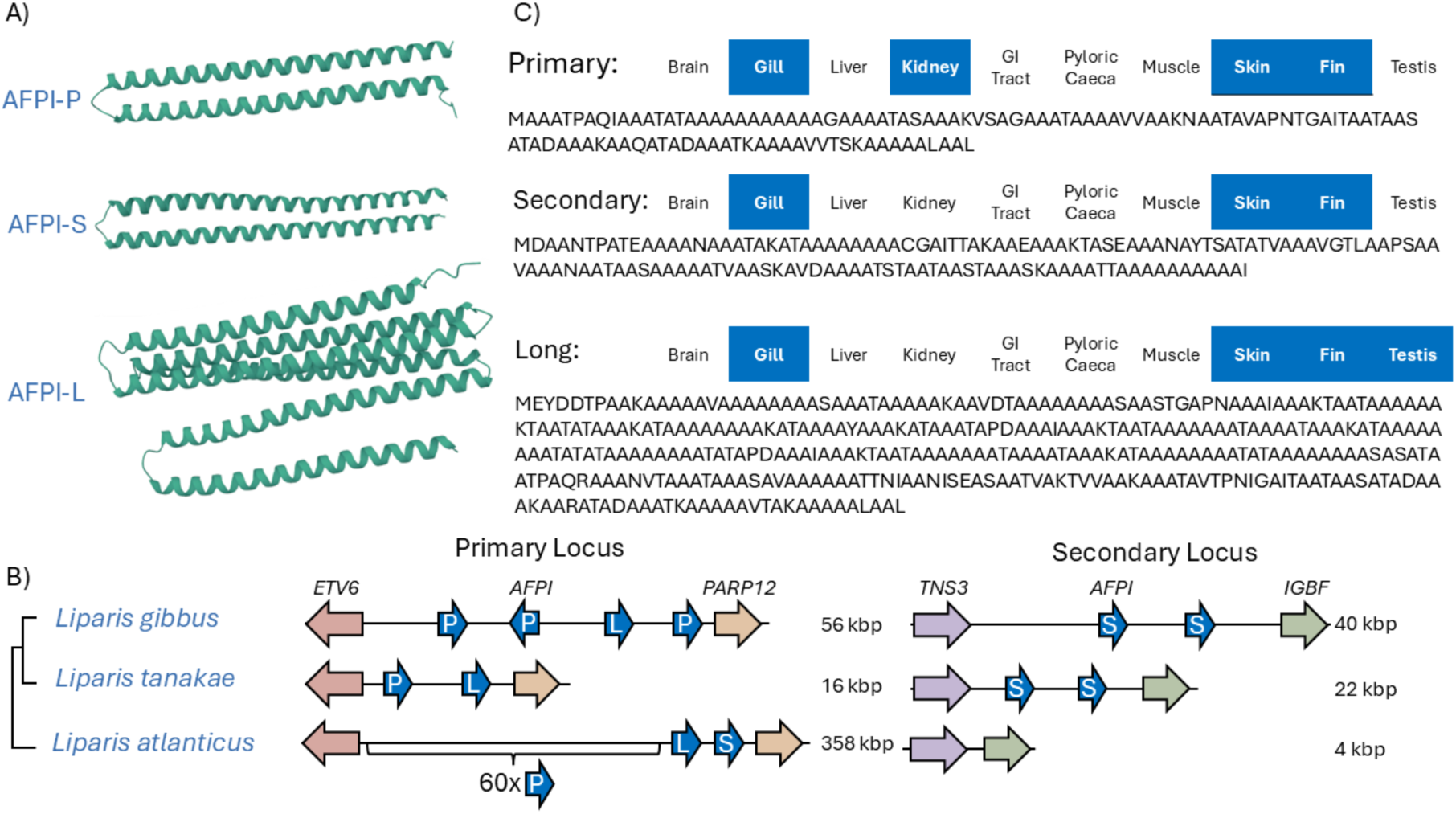
Structure, sequence, expression, and genomic positions of snailfish AFPI isoforms. (A) Representative AlphaFold2 models of the three AFPI isoform classes, primary (AFPI-P), secondary (AFPI-S), and long (AFPI-L), showing predominantly alpha-helical structures. (B) Locus maps showing the genomic positions of AFPI isoforms in three AFPI-bearing Liparis species. Colored arrows indicate genes and their transcriptional orientation; arrow length is not to scale. P, S, and L denote primary, secondary, and long isoforms, respectively, and locus sizes indicate the distance between flanking genes. (C) Representative amino acid sequences of each isoform class, illustrating differences in terminal motifs and in the extent of the alanine-rich repetitive tract; colored boxes indicate tissues in which each isoform was detected by RNA-seq.

The secondary isoform is intermediate in length between the long and primary isoforms. Where a secondary locus is present, secondary isoforms are associated with that locus; however, *L. atlanticus*, which lacks a secondary locus, still contains a secondary isoform located in the primary locus (Fig. 2B).

The three isoform classes also show characteristic terminal motifs (Fig. 2C). Primary isoforms begin with MA[P/A]ATP and typically end with leucine, often as LAAL. Secondary isoforms begin with MDAANT and end with isoleucine. Long isoforms begin with MEY[D/Q]DTP or MDAATP and terminate with either isoleucine or leucine, depending on species. Compared with AFPI sequences from sculpin, flounder, and cunner examined in our previous work [38], snailfish AFPI sequences are more heterogeneous and contain less regular alanine-rich repeat structure. Predicted protein structure remains predominantly alpha-helical, but includes helix-breaking residues not reported in other AFPI lineages. Overall, snailfish AFPI exhibits the greatest diversity in copy number, sequence architecture, and interspecific variation among the AFPI lineages examined.

We next examined isoform-specific expression across the ten tissues sampled for RNA-seq (Fig. 2C). Previous work reported strongest AFPI expression in skin, with lower expression in liver [33]. In our data, all three isoform classes were expressed in gill, skin, and fin. Additional expression differences were isoform-specific: the primary isoform was also expressed in kidney, whereas the long isoform showed additional expression in testis. These results indicate that, although all AFPI isoforms are expressed in peripheral tissues associated with antifreeze function, individual isoform classes have diverged in tissue distribution.

### De novo origin of the alanine-rich AFPI coding sequence

To investigate the genomic context and potential origins of the AFPI coding sequence in snailfishes, we conducted genome-wide sequence-similarity searches using Iso-Seq–derived AFPI isoforms generated in this study, together with species-specific AFPI paralogs. BLASTN searches against the NCBI nt database, each AFPI+ snailfish genome, and closely related AFPI– genomes revealed no clear one-to-one homology between AFPI and any single protein-coding precursor. Within snailfish genomes, the only coherent locus-level matches were the AFPI-containing regions themselves; all other hits were isolated repeats or intergenic fragments. Comparative analyses of syntenic regions in AFPI– species further showed that the genomic interval corresponding to the AFPI coding region is either noncoding or lacks the long alanine-rich ORF, indicating that this tract is a derived feature of the AFPI+ snailfish lineage.

Because the AFPI coding region is composed largely of GC-rich tandem repeats encoding alanine-rich sequence, such a tract could readily have arisen by local duplication and expansion of a short, low-complexity noncoding motif. In this scenario, the ancestral proto-repeat in outgroup genomes would be both short and compositionally simple, making it difficult to distinguish from the genomic background of similar low-complexity sequences and helping to explain why clear orthologous noncoding sequences cannot be identified. This pattern is not unique to snailfish AFPI: similarly low-complexity, repetitive coding tracts evolved independently in the other three AFPI lineages and in the northern gadid AFGP gene [26, 38], suggesting that such sequence architecture may be especially prone to de novo emergence from noncoding DNA.

Searches against curated transposable-element libraries (Dfam) returned only low-confidence matches between short, compositionally biased segments of the AFPI coding region and gag-like domains of Copia long terminal repeat (LTR) retroelements. In *L. atlanticus*, two fragmented Copia-related elements occur within the AFPI locus, but both lack intact LTRs and the integrase domain, and homologous Copia elements elsewhere in the genome show no sequence similarity to AFPI. Related fragmented Copia-like sequences are also present in *L. gibbus* and *L. tanakae*, but again provide no evidence of a direct coding-sequence origin. We therefore find no support for wholesale co-option of a TE open reading frame into the alanine-rich core of AFPI. Instead, the limited similarity to TE sequence is best explained by convergent low-complexity repeat composition, consistent with the interpretation that the AFPI coding region evolved de novo from noncoding sequence and was subsequently elaborated through repeat expansion.

Among the low-complexity matches recovered in our genome-wide searches, the only gene showing notable similarity was an alanine-rich tract within exon 3 of *PARP12*, which lies adjacent to AFPI in the primary locus. This exon 3 tract is present in AFPI+ snailfishes but is absent from homologous *PARP12* exons in other teleosts. At the nucleotide level, the similarity between this *PARP12* region and AFPI is dominated by GCN codons encoding alanine. When positions corresponding to these alanine codons are ignored, identity drops sharply and no extended higher-complexity matches remain. Thus, although *PARP12* exon 3 and AFPI share a locally expanded alanine-rich repeat, the observed similarity does not establish direct homology or a clear direction of origin.

Genome-wide comparisons also identified additional *PARP12*-related sequences associated with the AFPI loci. Outside snailfishes, a *PARP12*-like gene containing the *PARP12* first three exons occurs in multiple teleosts, including as a functional protein-coding gene in ballan wrasse and channel bull blenny and as a long noncoding RNA in stickleback (Supplementary Fig. 3), but this gene is absent from all examined snailfish genomes, including both AFPI+ and AFPI− species. In AFPI+ snailfishes, however, fragments related to the broader *PARP12* region, most notably exon 1 and the adjacent intronic sequence, are repeatedly duplicated alongside AFPI genes within both AFPI loci. Together, these observations indicate that *PARP12*-related sequence has been repeatedly copied and remodeled in this genomic neighborhood, but neither the intact *PARP12* gene nor the *PARP12*-like gene appears to have served as a direct progenitor of the AFPI coding sequence.

### Contribution of TE fragments to AFPI regulatory sequences

Building on the evidence that the alanine-rich AFPI coding sequence likely arose from previously noncoding or minimally coding sequence, we next characterized the origins of its untranslated regions (UTRs) and adjacent flanking sequences, as they are potential regulatory sequences.

The beginning of the 5′ UTR and the immediately upstream flanking region share high sequence identity with a hAT-family DNA transposon (Fig. 3), indicating that the AFPI transcription start region and cis-regulatory sequence were derived from a transposon insertion. This retained hAT-derived fragment contains two putative core promoter motifs, a BRE and a TATA box, consistent with a role in transcriptional regulation. Supporting this interpretation, similar hAT-derived sequences are conservatively found in intronic or flanking regions of other protein-coding genes in snailfishes and other species, including *copine-3*-*like*, *NLRX1*, and *ZBED1-like*. To infer the origin of this regulatory sequence, we examined the AFPI-associated hAT fragment. The sequence most closely matches a non-autonomous hAT-like element from zebrafish, as it lacks transposase coding domains. Additionally, the fragment lacks the terminal inverted repeats (TIRs) required for mobilization, pointing to post-insertion sequence decay. Together, these structural features indicate that the AFPI 5′ regulatory region preserves a degraded remnant of an ancestral hAT-derived insertion. The retained sequence, however, still contains promoter-like motifs, consistent with subsequent co-option of hAT-derived sequence for AFPI transcription.

**Figure 3.**
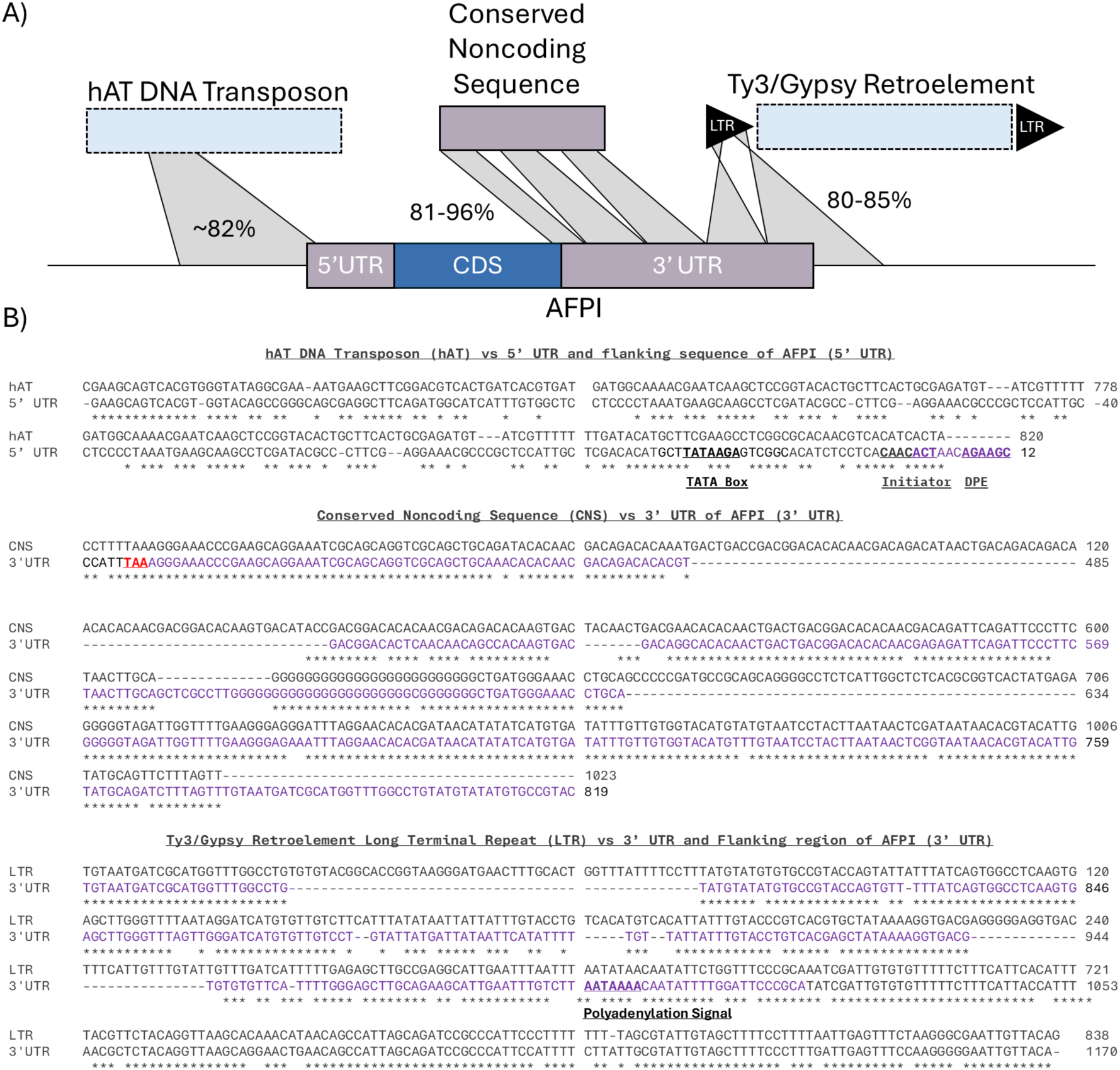
Sequence architecture of AFPI regulatory regions. (A) Schematic summary of sequence segments homologous to the AFPI gene and flanking sequences. The 5’ untranslated regions (UTR) and immediately upstream flanking region are homologous to a hAT-family DNA transposon, the sequence spanning the terminal coding nucleotides, stop codon, and proximal 3’ UTR is homologous to a conserved noncoding sequence, and the distal 3’ UTR together with downstream flanking sequence is homologous to the long terminal repeat (LTR) of a Ty3/Gypsy retrotransposon. Numbers indicate the range of nucleotide identity between AFPI and each homologous source region. (B) Nucleotide alignments between the hAT element and the AFPI 5’ UTR plus upstream flanking sequence; between the conserved noncoding sequence and the 3’ end of AFPI; and between the Ty3/Gypsy LTR and the distal 3’ UTR plus downstream flanking sequence of AFPI.

The distal portion of the AFPI 3′ UTR shares strong sequence similarity with the LTR of a Ty3/Gypsy retrotransposon (Fig. 3), suggesting that part of the downstream AFPI regulatory region was derived from an LTR fragment. Because LTR retrotransposons use their LTRs in transcript-end formation, including cleavage and polyadenylation, co-option of such a fragment into a host 3′ regulatory region is mechanistically plausible [44]. Although the source retrotransposon is now fragmented at the AFPI loci, homologous Ty3/Gypsy-like retroelements are still detectable in all three AFPI+ species. The persistence of the same identifiable LTR-derived segment across all AFPI genes, despite broader retroelement fragmentation in the locus, points to functional retention after incorporation into the AFPI 3′ UTR. These observations support the interpretation that the AFPI 3′ UTR retains a degraded remnant of an ancestral Ty3/Gypsy insertion that became incorporated into the downstream regulatory region.

Adjacent to the LTR-derived distal segment, the proximal portion of the 3′ UTR, together with the stop-codon region and the terminal few nucleotides of the coding sequence, shares high identity with a deeply conserved noncoding sequence (Fig. 3). Its conservation across fishes is supported by the Ensembl 65-fish EPO-Extended whole-genome alignment [45], consistent with functional constraint. In the primary AFPI locus, the homologous sequence lies immediately upstream of the transcription start site of *ETV6*, the gene adjacent to AFPI, indicating a highly conserved proximal regulatory region. In AFPI+ snailfish, the AFPI-homologous portion of this conserved noncoding block, together with its adjacent flanking sequence, is locally expanded relative to AFPI− outgroups. Together, these observations suggest that local duplication or repeat expansion of this conserved noncoding block preceded AFPI origination and may have supplied sequence later incorporated into the 3′ end of the AFPI transcript. The sequences abutting this conserved block are also highly repetitive and GC rich, making them plausible noncoding substrates for the repeat expansion that accompanied de novo origination of the alanine-rich, GCN-encoded AFPI coding sequence.

### TE-associated mechanisms for AFPI translocation and copy-number change

Repeat annotation showed that both primary and secondary AFPI loci in snailfishes are embedded in dense landscapes of interspersed repeats. Most annotated repeats correspond to fragmented TEs, although a subset of intact elements remains. Across AFPI+ species, AFPI genes occur within expanded, TE-rich intervals whose size ranges from a few kilobases to several hundred kilobases (Fig. 4). For example, the primary AFPI locus in *L. atlanticus* spans over 300 kb and contains 62 AFPI genes, whereas the primary locus in *L. tanakae* has contracted to less than 10 kb and no longer contains AFPI in some populations. These observations indicate that AFPI resides in highly dynamic genomic neighborhoods where TE insertion, gene duplication, and deletion have contributed to rapid locus remodeling. The extensive accumulation of TE-derived sequence around AFPI, together with marked variation in AFPI copy number and the occurrence of AFPI at two chromosomal loci, points to repeat-mediated mechanisms as plausible drivers of both locus movement and family expansion.

**Figure 4.**
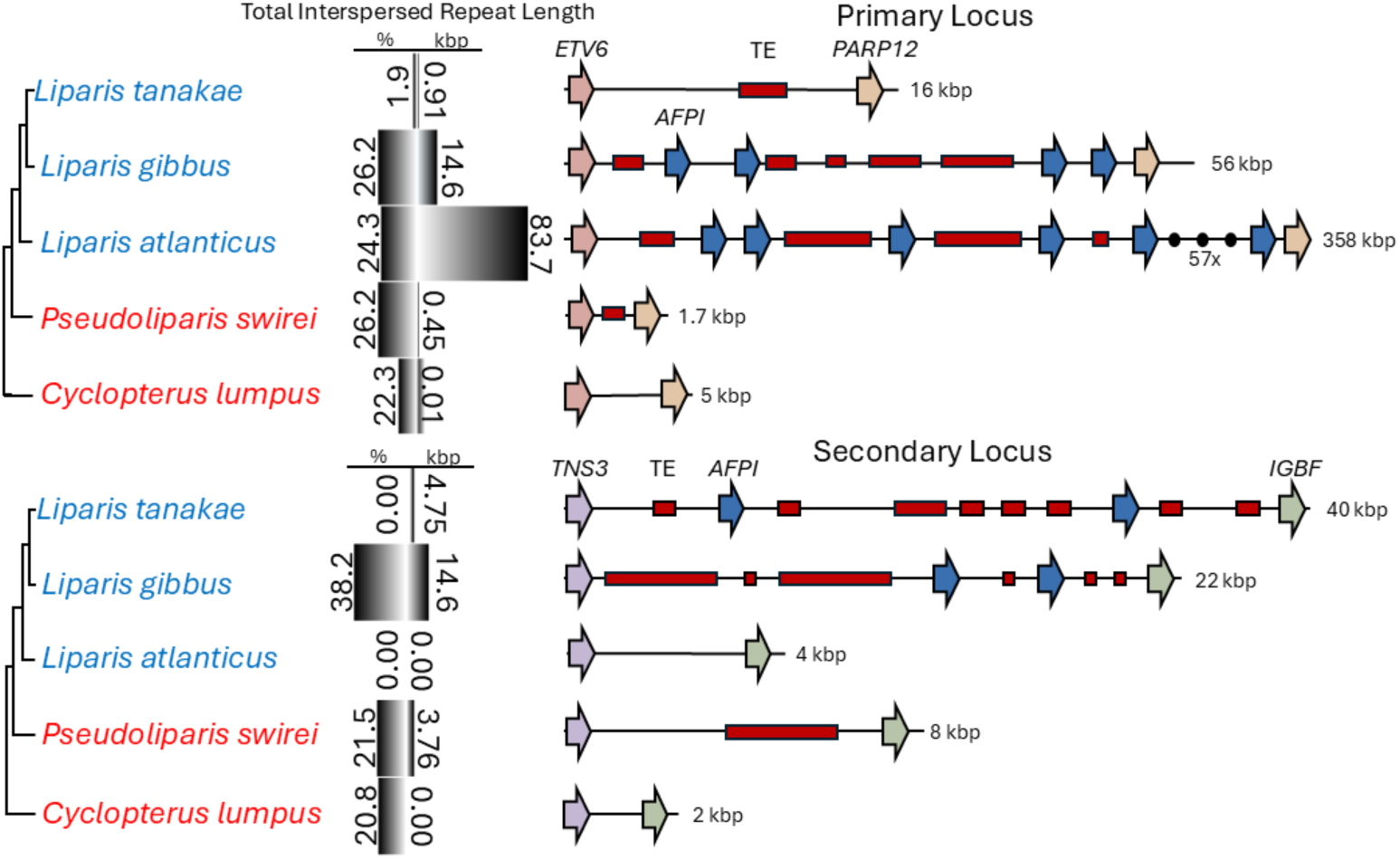
Transposable element content and repeat architecture of AFPI loci in AFPI-bearing and AFPI-lacking species. Total interspersed repeat content is shown within the primary AFPI locus and the secondary locus. Values are shown as the percentage of each locus annotated as interspersed repeats (left side) and the corresponding repeat length in kilobases (right side). Repeat content was calculated across the interval between the two flanking genes defining each locus. The locus maps show the distribution of repeat annotations across the primary and secondary AFPI loci.

One question is how an AFPI-bearing segment reached the secondary locus. Given the inferred Miocene origin of AFPI within *Liparis* and the substantial subsequent remodeling of these loci, the absence of fully intact transposons or complete insertion signatures is not unexpected. Even so, the remaining sequence architecture implicates TE-associated movement. In *L. tanakae*, TIR-associated sequence flanks AFPI genes, and in *L. gibbus* a fragmented mariner-like element occurs adjacent to AFPI-linked sequence. Tc1/mariner elements are cut-and-paste DNA [46, 47] transposons whose transposases act on DNA delimited by terminal inverted, so these features support a DNA transposon-associated mobilization model for transfer of an AFPI-containing segment (Fig. 5A). Homologous LINE fragments shared between the primary and secondary loci of *L. gibbus* provide a second, independent line of evidence. In vertebrates, LINE-1 elements can mobilize downstream host DNA by 3′ transduction and thereby copy adjacent genomic sequence to new loci [48]. Taken together, the presence of TIR-associated sequence, mariner-like fragments, homologous LINEs, and shared AFPI-flanking sequence between loci supports TE-associated acquisition of the secondary locus rather than independent origin of AFPI at that site (Fig. 5B).

**Figure 5.**
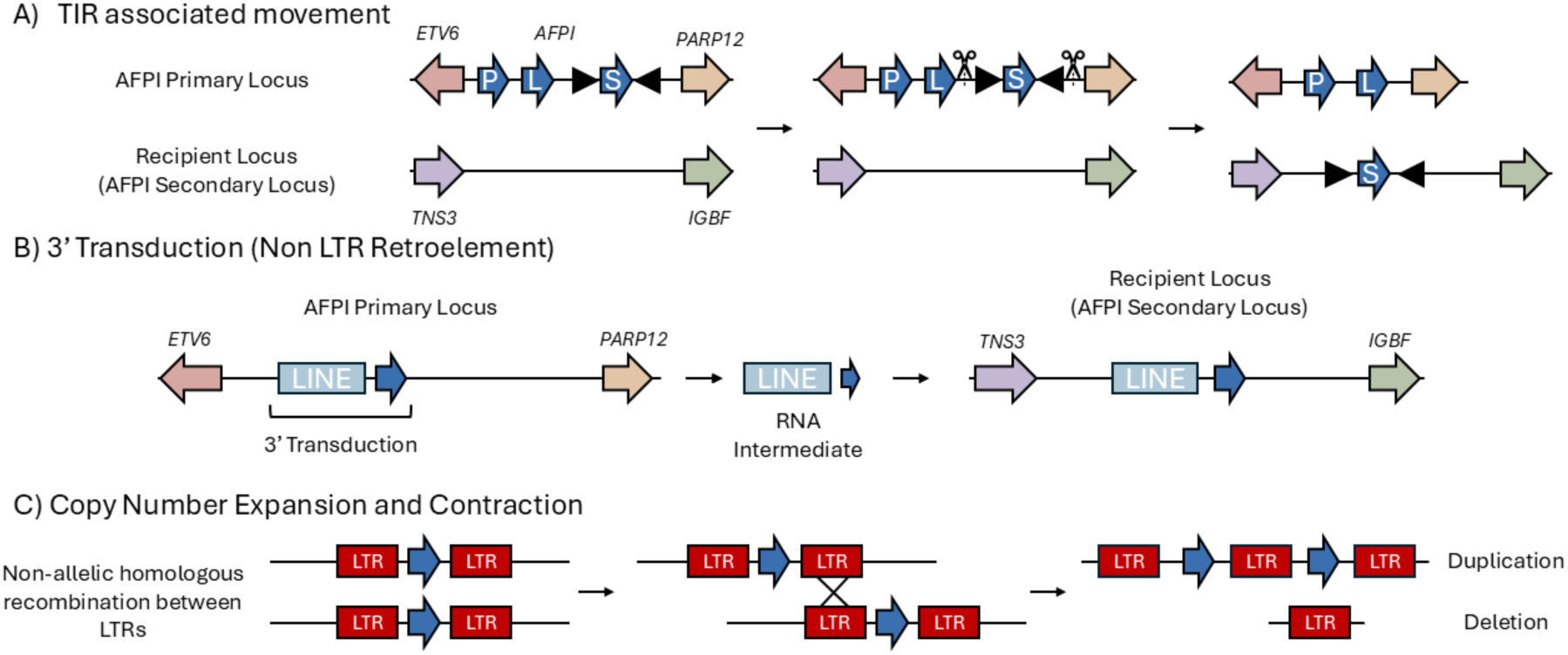
Proposed TE-associated mechanisms for AFPI translocation and copy-number changes. (A) DNA transposon-mediated movement of an AFPI-containing DNA segment, motivated by the presence of TIR-associated sequence in AFPI loci. (B) LINE-associated 3’ transduction, in which a LINE transcript reads through into adjacent AFPI sequence and copies downstream host DNA to a new locus. (C) Proposed mechanism for AFPI copy-number change within loci. Non-allelic homologous recombination between misaligned long terminal repeat (LTR)-derived repeats could generate reciprocal duplication and deletion products.

Within loci, LTR-derived repeat structure provides the clearest mechanistic explanation for AFPI copy-number change. Multiple homologous LTR retroelements from several TE families, including Gypsy and Copia, were found exclusively in AFPI+ species. Because non-allelic homologous recombination between repeated elements is a well-established source of reciprocal duplication and deletion [49], these shared LTR-derived segments provide a strong substrate for AFPI copy-number expansion and contraction. This mechanism parsimoniously explains the extensive variation in AFPI copy number observed among AFPI+ species and among populations of *L. tanakae* (Fig. 5C).

## Discussion

### Convergent AFPI evolution across four lineages reveals independent molecular routes to partial de novo gene formation

This study clarifies the origin and mechanism of the new gene, AFPI, in the fourth AFPI+ lineage, completing the account of AFPI new gene origination across all four AFPI+ lineages. Across these distantly related taxa, AFPI has arisen independently yet convergently, not only in function but also in protein architecture. Facing similar freezing-selection pressures associated with recent marine glaciations (polar or subpolar conditions) [50], each lineage independently acquired a new gene with the same adaptive function, enabling survival in lethal polar waters. In this rare case of protein convergent evolution, selection repeatedly favored short alanine-rich repeats with regularly spaced threonine that confer ice-binding activity [36]. Despite divergent nucleotide sequences and codon usage, these proteins converge on similar α-helical structures for ice recognition [51].

Beneath this striking convergence, however, the genetic routes by which AFPI originated differ sharply among lineages. In our previous work, we showed that in the flounder lineage AFPI arose by recruiting noncoding portions of the antiviral gene GIG2; in cunner, AFPI originated via gene fission of the immune-regulatory GTPase GIMAP4; and in sculpin, AFPI assembled by incorporating fragments of a pseudogene derived from the ER network-shaping protein LNPK-B [38]. In the snailfish lineage examined here, AFPI instead derives from a mosaic of TEs and conserved noncoding sequences. Although the mechanisms are unique in each case, all four lineages share a common theme of partial de novo evolution, in which most of the coding sequence arose de novo from noncoding sequence, whereas regulatory sequences were recruited and modified from pre-existing protein-coding precursors or TEs. Viewed together, AFPI now represents four independent molecular routes to the same adaptive protein, making it an exceptional example of convergent evolution at both the functional and protein sequence levels.

### Snailfish AFPI extends the Duplication-Degeneration-Divergence model of new gene origination

In our previous study, we proposed the Duplication-Degeneration-Divergence (DDD) model to explain how degenerated functional units can be reorganized into novel genes with unrelated functions [38]. The snailfish lineage extends this framework in an important way. Unlike the other three AFPI-bearing lineages, in which AFPI traces back to a single precursor gene, snailfish AFPI appears to have arisen from multiple genomic sources, including different TEs and noncoding sequences. In this case, degeneration, fragment recruitment, and sequence reorganization did not simply modify an existing gene or split ancestral functions among duplicates. Instead, these processes contributed to the assembly of a novel protein-coding gene. The snailfish case therefore broadens the DDD model by showing how genomic fragments that no longer function in their original context can be repurposed into a new adaptive gene.

More broadly, the snailfish case highlights how “genomic debris” can contribute to evolutionary innovation. Rather than beginning from a single ancestral coding template, snailfish AFPI appears to have emerged from a genomic neighborhood in which decayed functional fragments, conserved noncoding sequence, and TE-derived regulatory elements were brought into proximity and then reorganized. Once an incipient gene product with ice-binding potential emerged, strong environmental selection likely amplified and refined it. AFPI therefore offers an exceptional system for dissecting how novel proteins can evolve through multiple genetic routes under a shared selective regime. The convergence of sequence features and biochemical function despite mechanistically distinct origins highlights a general principle: strong, predictable selection can repeatedly sculpt very different genomic substrates into similar adaptive solutions.

### Transposable elements in new gene origination and diversification

Our results extend the well-established role of TEs in evolutionary innovation by documenting a case in which TE-derived sequence is associated with the assembly and diversification of a new gene, rather than merely modulation of an existing locus. Within *Liparis*, the AFPI-bearing loci are strongly enriched for TE-derived sequence without a comparable expansion at orthologous loci in closely related AFPI− taxa. In snailfish AFPI, TE signatures are linked to multiple stages of gene evolution, including a hAT-derived fragment at the 5′ end associated with promoter-like motifs, a Ty3/Gypsy-derived fragment in the distal 3′ UTR, and repeat-rich local architecture that provides plausible substrates for duplication and translocation. Together, these observations place TEs at the center of the molecular changes that turned a previously noncoding genomic neighborhood into a functional, rapidly diversifying antifreeze locus.

This interpretation fits within a broader literature in which TEs generate novelty through several routes, including domestication of TE coding sequence, exaptation of TE-derived fragments, RNA-mediated retrogene formation, and especially rewiring of gene regulation through promoter and enhancer recruitment [10, 52]. However, the best-characterized regulatory examples generally involve pre-existing genes. In sablefish, for example, a hAT insertion in the *gsdfY* promoter brought cis-regulatory modules that rewired expression of a duplicated gene [53], and in black carp TE-derived promoter and enhancer elements promoted dmrt1 diversification into a male-determining locus [54]. More relevant to de novo evolution, DNA transposons in Drosophila are associated with the emergence of new transcripts by contributing upstream transcription factor-binding motifs [19], and in roses a MITE insertion boosts promoter activity of the de novo-originated *SCREP* gene [55]. Snailfish AFPI goes a step further: the data support TE contribution to both ends of the transcription unit and to the structural dynamics of the locus itself. In this system, TE-derived sequence appears to provide parts of the transcriptional framework, including the 5′ regulatory region and distal 3′ regulatory sequence, while TE-rich repeats furnish mechanisms for duplication and locus movement. In that sense, snailfish AFPI is not just a case of TE-mediated expression change. It is a mechanistically explicit example of how TE-derived sequence can help convert noncoding DNA into a functional vertebrate gene and then promote its subsequent expansion and diversification.

The evolutionary role of TEs in AFPI does not appear to stop at gene origination. TE-associated processes also provide plausible mechanisms for the subsequent duplication and movement of AFPI after the gene was established. TE-mediated duplication and relocation of host genomic material are well documented, including through recombination between repetitive elements and LINE-associated transduction of flanking DNA [56–58]. However, these mechanisms have usually been discussed in the context of duplicating or reshuffling pre-existing genes or gene fragments [56, 59]. In snailfish AFPI, by contrast, TE-associated processes appear to act on a locus whose coding sequence most likely emerged de novo from noncoding sequence, linking TE-facilitated locus movement to the assembly and diversification of a newly evolved gene.

### TE-rich AFPI loci as localized substrates for innovation under freezing selection

Although TEs likely contributed directly to AFPI assembly, duplication, and translocation, their influence may also have depended on the broader character of the surrounding locus. In AFPI+ *Liparis* species, AFPI loci are strongly enriched for TE-derived sequence and show signatures of duplication, translocation, and fragment recruitment, whereas orthologous regions in AFPI-lacking snailfish lineages show little comparable expansion or remodeling. This contrast argues against a simple genome-wide TE effect and instead points to a localized genomic neighborhood that was especially permissive to innovation.

TE activity is known to fluctuate between relative quiescence and episodic bursts, and TE-dense genomic “islands” can accumulate structural variants and copy-number changes during ecological transitions [16, 60]. In the context of progressive ocean cooling and glaciation, AFPI loci in *Liparis* may represent a similar class of repeat-rich regions in which regulatory fragments, low-complexity sequence, and recombination substrates were repeatedly brought together and remodeled under strong freezing selection. In this view, the key contribution of TEs in the AFPI system is not simply higher repeat content, but the creation of a local genomic environment that favored new gene emergence and rapid family-level diversification.

In summary, snailfish AFPI completes the comparative picture of AFPI evolution by revealing a fourth, mechanistically distinct route to the same adaptive protein. It shows that a young vertebrate gene family can arise de novo from noncoding or minimally coding DNA, while TE-derived fragments and TE-rich locus dynamics contribute regulatory sequence, duplication, and translocation. More broadly, this case shows that convergent adaptation can be achieved from very different genomic substrates, and that TEs can participate not only in altering gene expression, but also in the assembly, expansion, and remodeling of a newly evolved vertebrate gene. AFPI therefore provides an unusually clear model that illustrates how strong selection can repeatedly shape very different genomic substrates into similar adaptive outcomes.

## Materials and Methods

### Specimen Collection and Dissection

Adults of Atlantic snailfish (*Liparis atlanticus*) were collected from Cobscook Bay near Pembroke, Maine, in spring 2023 by Gulf of Maine Inc. Specimens were anesthetized with tricaine methanesulfonate (MS-222) and placed on ice for blood collection via caudal venipuncture. While maintained on ice, tissues including brain, gill, liver, kidney, gastrointestinal tract, pyloric caeca, muscle, skin, fin, and testis/ovaries were dissected. All tissues were fresh frozen in liquid nitrogen immediately after dissection and stored at −80 °C until use. The handling and sampling of fish complied with protocol #23026 approved by the University of Arkansas Institutional Animal Care and Use Committee (IACUC).

### Long-Read Genome Sequencing and Assembly

Blood cells from one individual specimen were used in whole genome sequencing. High molecular weight (HMW) DNA was extracted from blood cells using the Nanobind CBB kit (PacBio) following the manufacturer’s protocol optimized for extracting HMW DNA from nucleated red blood cells. The HMW DNA sample was sequenced on a PacBio Revio System using one SMRT Cell 25M with a 30-hour movie, performed by the Genome Technology Access Center at the McDonnell Genome Institute, Washington University School of Medicine. PacBio HiFi circular consensus sequencing (CCS) generated 3.14 million high-accuracy reads totaling 60.2 Gbp, approximately 102× coverage of our assembled genome and ∼82× coverage based on the estimated genome size for other *Liparis* genus species reported in the Animal Genome Size Database (http://www.genomesize.com).

PacBio HiFi reads were assembled de novo using hifiasm V. 0.19.5 [61] with default parameters to generate a primary contig-level assembly. Assembly completeness was evaluated using BUSCO V. 5.8.3 [62] with the Actinopterygii reference gene set, which indicated high completeness (96.4%) but an elevated proportion of duplicated BUSCO genes (64%), consistent with residual haplotigs. To reduce redundancy, haplotigs and overlapping contigs were identified and removed using purge_dups V. 1.2.6 [63] with default settings, and assembly quality was reassessed with BUSCO, which showed that the proportion of duplicated genes decreased to 1.8% after purging.

### Hi-C Sequencing and Genome Scaffolding

To scaffold the draft genome assembly to chromosome scale, Hi-C library preparation and sequencing was performed by Phase Genomics (Seattle, WA) using muscle tissue from *L. atlanticus*. Sequencing was carried out on an Illumina NovaSeq X Plus platform, generating 184.3 million paired-end reads (2 × 150 bp). The Hi-C based contig scaffolding was performed using SALSA2 v. 2.3 [64] Hi-C reads were aligned to the draft genome assembly using the juicer pipeline [65] to generate contact maps including the BWA v. 0.7.17 [66] dependency. Chromosome-scale scaffolding was subsequently refined using both the 3D-DNA pipeline [67] and YaHS v. 1.2.2 [68]. Assemblies generated by these approaches were evaluated using BUSCO V. 5.8.3; while completeness metrics were comparable, the YaHS-scaffolded assembly exhibited greater contiguity and was retained for downstream analyses.

### RNA Isolation and Iso-Seq Transcriptome Sequencing

To generate full-length transcript evidence for genome annotation, total RNA was isolated from ten tissues of one *L. atlanticus* male adult (brain, gill, liver, kidney, gastrointestinal tract, pyloric caeca, muscle, skin, fin, and testis). Approximately 20–25 mg of each tissue was homogenized in TRIzol reagent (Invitrogen), and total RNA was extracted following the manufacturer’s protocol. Residual genomic DNA was removed by DNase I treatment (New England Biolabs), and RNA was purified using the Monarch RNA Cleanup Kit (New England Biolabs). RNA quantity and purity were assessed using Qubit and NanoDrop assays, and RNA integrity was evaluated with an Agilent TapeStation; all samples exhibited high integrity (RIN > 8). Purified RNA from each tissue was pooled in equimolar amounts (4 μg total RNA) to maximize transcript diversity for Iso-Seq library preparation. Iso-Seq library construction (Poly(A) enrichment; PacBio Kinnex) and sequencing were performed by Novogene Corporation Inc. (Sacramento, CA) on a PacBio Revio platform, generating approximately 10 million raw reads.

IsoSeq reads were processed using the PacBio tool suite (https://github.com/PacificBiosciences/pbbioconda). Lima v. 2.13.0 was first used with default settings to demultiplex based on barcode sequences to assign reads to their corresponding libraries. The isoseq v. 4.3.0 package was used to identify full-length non-chimeric (FLNC) reads with command isoseq refine using the flags --require-polya and --min-polya-length 20 to ensure transcript completeness. Refined reads were clustered and polished with the isoseq cluster function to generate high-quality consensus transcripts, which were mapped to the genome using Minimap2 v. 2.29 [69].

### Tissue-specific Short-read RNA-Seq and Expression Analysis

To examine tissue-specific expression patterns of AFPI isoforms and related genetic components, short-read RNA-seq data were generated and analyzed across ten tissues. Purified total RNA from the ten tissues (listed above) was used for Illumina library construction and RNA sequencing at the Clinical Genomics Center, Oklahoma Medical Research Foundation. Libraries were sequenced on an Illumina NovaSeq X Plus platform to a depth of approximately 30 million paired-end reads (2 × 150 bp) per tissue.

RNA-seq reads were aligned to the reference genome using STAR v2.7.11b [70], and gene-level read counts were obtained with featureCounts (Subread package) [71] using gene models generated by BRAKER2 v. 3.0.8 [72–75]. Raw counts were normalized using the variance-stabilizing transformation (VST) implemented in DESeq2 [76] to facilitate comparison of expression patterns across tissues. Normalized expression values were compared descriptively across tissues to identify tissue-associated expression patterns, with particular focus on AFPI isoforms. For each AFPI isoform, expression status (presence/absence) was assessed across tissue types and visualized in R using ggplot2 [77].

### Genome Annotation

Genome annotation was performed on a soft-masked assembly using BRAKER2 v. 3.0.8, with protein homology. The genome was softmasked using RepeatMasker v. 4.2.2 [78] with the xsmall command to return repeats in lowercase rather than complete masking. Full-length transcript models generated from PacBio Iso-Seq data were incorporated to improve exon–intron structure, alternative splice isoform, and UTR annotations. Protein evidence supplied to BRAKER2 included curated vertebrate protein sequences from OrthoDB [79] as well as annotated proteomes from the closely related species *Pseudoliparis swirei* and *Cyclopterus lumpus*. This integrative annotation strategy was used to generate gene models for downstream comparative and evolutionary analyses.

### Genome-Wide Transposable Element Annotation

To characterize genome-wide TE content, RepeatModeler v. 2.0.7 [80] was run independently on each genome assembly to generate de novo repeat libraries. Species-specific libraries were combined with the broad Dfam RepeatMasker library [81] to create a comprehensive repeat library capturing both conserved and lineage-specific elements. This combined library was used to soft-mask each genome using RepeatMasker, producing standardized summary statistics of TE content across species (Supplementary Fig. 1).

### Locus-Specific Transposable Element Characterization at AFPI Loci

To enable in-depth identification of intact and fragmented TEs associated with AFPI loci, we applied a complementary set of similarity-based, structure-based, and de novo approaches, which are necessary for non-model organisms lacking curated TE libraries. Initial similarity searches were conducted at AFPI-associated genomic intervals using RepeatMasker with Dfam profiles, including teleost-derived TE profiles (e.g., zebrafish TE libraries). This approach was used to identify candidate TE-derived sequences and fragments without assuming complete element structure. Candidate nucleotide sequences were subsequently screened against the NCBI Conserved Domain Database (CDD) to detect TE-associated domains (e.g., reverse transcriptase, integrase). When intact open reading frames were identified, predicted protein sequences were further analyzed using Pfam [82], PROSITE [83], and NCBI-CDD [84] for domain characterization. Additionally, FISHTEDB [85] was queried to identify fish-specific elements. Structural TE annotation was further supported using EDTA [86], which integrates LTR_retriever, TIR-Learner, and HelitronScanner. Evidence from similarity-based, structure-based, and de novo approaches was integrated and manually curated at AFPI loci in Fig. 4.

### Characterization of AFPI Gene Family

To identify AFPI genes across all available genomes, AFPI loci were manually annotated using a homology-based approach. Publicly available AFPI cDNA and protein sequences from GenBank for *Liparis gibbus* (AY455863.2) and *Liparis atlanticus* (AY455862.2) were used as queries in genome-wide BLASTN and BLASTP searches against all five snailfish genomes using the BLAST+ suite [87]. Searches were conducted without low-complexity filtering due to the repetitive nature of AFPI coding sequences. Additional searches using the non-repetitive flanking regions and UTRs were performed to identify AFPI copies regardless of isoform or repeat length. Candidate loci were inspected and manually annotated using SnapGene (www.snapgene.com), and complete AFPI gene sequences were reconstructed. Isoform structures were annotated using full-length transcripts generated from Iso-Seq data. A representative amino acid sequence for each isoform was then used as input for ColabFold to predict protein structure, using default settings with msa_mode set to single_sequence [88]. The resulting structures were visualized using Mol* Viewer [89].

### Characterization of AFPI Genomic Loci in AFPI+ Species and Syntenic Region in AFPI– Species

To define the genomic loci harboring AFPI genes, scaffolds containing AFPI in each AFPI+ species were examined to identify flanking genes and intergenic regions. BRAKER2 annotations for *L. atlanticus* were imported into SnapGene for visualization. Protein-coding genes flanking AFPI loci were validated using BLASTP searches with default parameters to confirm gene ID and accuracy of the annotation (i.e. correct reading frame, start and stop codon, intron/exon junctions). If the protein sequence was unavailable, BLASTN searches were used to identify homologous nucleotide sequences of known genes. For species lacking available genome annotations, gene identity was inferred using annotations from the closest related species based on sequence similarity and conserved domain structure.

To identify syntenic genomic regions corresponding to the AFPI locus in AFPI– species, flanking genes and intergenic sequences from AFPI+ species were used as anchors. Orthologs of flanking genes were identified using BLASTP or BLASTN searches described above.

Annotation was guided by manual curation and by existing annotations from *Pseudoliparis swirei* and *Cyclopterus lumpus*, enabling direct comparison of locus structure across AFPI+ and AFPI– species. Additionally, the AFPI-containing scaffold in the *L. atlanticus* (AFPI+) genome assembly and the chromosome harboring the syntenic region in *Pseudoliparis swirei* (AFPI–) were aligned using minimap2, and the resulting alignments were visualized as a synteny plot with NGenomeSyn [90] (Supplementary Fig. 4).

### Inference of the Origins of the AFPI Coding and Regulatory Sequence

To test whether the alanine-rich AFPI coding region was derived from a pre-existing gene or from previously noncoding sequence, we first used mapped Iso-Seq reads to define full-length AFPI transcript models in *Liparis atlanticus* and combined these with annotated AFPI paralogs from the three AFPI+ species. Representative full-length transcripts from each AFPI isoform class, together with species-specific AFPI copies, were used as BLASTN queries in genome-wide searches against the NCBI nt database, the three AFPI+ genomes, and three AFPI− relatives/outgroups. Genome-wide rather than locus-restricted searches were used because the ancestral source sequence might not be retained within the present AFPI locus. Low-complexity filtering was disabled because the AFPI coding region is dominated by repetitive alanine-rich sequence. Hits recovered outside annotated AFPI loci were examined individually by additional searches against nt and the study genomes to determine whether they matched known protein-coding genes, noncoding regions, or low-complexity repeats. Candidate regions were then aligned to AFPI queries with MUSCLE [91] and manually inspected to assess alignment continuity and to exclude similarities driven primarily by short repetitive tracts.

We next separated the AFPI flanking and untranslated regions from the repetitive coding tract and performed additional BLASTN searches using the 5′ upstream region, 5′ UTR, 3′ UTR, and downstream flanking sequence as independent queries. All non-AFPI BLAST hits in snailfish genomes corresponded to noncoding sequence, including intergenic regions, UTRs, and introns of other genes. These sequences were subsequently queried against the Dfam and FishTEDB databases to evaluate whether they were derived from TEs. Candidate TE-derived and noncoding matches were compared with AFPI-associated sequence by MUSCLE alignment, and homology was evaluated by manual inspection of alignment length, continuity, and percent identity.

Putative core promoter motifs in the AFPI 5′ region were predicted with ElemeNT 2023 [92]. To evaluate whether AFPI-homologous noncoding segments corresponded to conserved regulatory sequence in outgroup teleosts, orthologous regions at the primary locus were further examined in Ensembl [45], and multispecies conservation was assessed using the Ensembl 65-fish EPO-Extended whole-genome alignment, with particular attention to AFPI-homologous sequence adjacent to *ETV6*.

### Phylogenetic Analyses of Snailfish Species and AFPI Genes

We reconstructed the phylogeny of the snailfish species in this study using *Cyclopterus lumpus* as the outgroup, employing two complementary approaches (Fig. 1). First, single-copy BUSCO orthologs (amino acid sequences) present in all taxa were extracted from genome assemblies of these species. These orthologs were concatenated and aligned using MAFFT v. 7.525 [93]. The best-fitting substitution model was selected using raxmlGUI 2.0 [94], and a maximum-likelihood phylogeny was inferred with RAxML-NG [95] under the JTT+I+G4+F model. Second, to provide an independent, DNA-sequence–based phylogenetic inference, we applied Read2Tree [96], which infers species trees directly from nucleotide sequence data using predefined orthologous markers. DNA sequence reads from *Pseudoliparis swirei* (SRX16006097), ,*Cyclopterus lumpus (VGP ID:* fCycLum2*), Liparis atlanticus*, *Liparis gibbus* (SRX24536308), and *Liparis tanakae* (SRX27879558) were analyzed with Read2Tree, using orthologous markers selected from the OMA (Orthologous Matrix) database [97] based on 100 DNA markers that covered at least 80% of teleost species in the database. Markers shared across all taxa were aligned and used to infer a species tree. The Read2Tree topology was concordant with the BUSCO-based amino-acid phylogeny, supporting the robustness of the inferred species relationships.

To reconstruct the evolutionary relationships among AFPI genes (Supplementary Fig. 2), all annotated AFPI gene sequences were aligned using MAFFT. Poorly aligned and spurious regions were removed using trimAl [98]. The optimal nucleotide substitution model was determined using ModelTest-NG [99], and a maximum-likelihood phylogeny was inferred with RAxML-NG under the GTR+G4 model.

A broader teleost species phylogeny (Supplementary Fig. 3) was obtained using the FishTree R package [100, 101], which draws from the Fish Tree of Life database. All phylogenetic trees were visualized and annotated using iTOL [102].

## Acknowledgments

We acknowledge the Arkansas High Performance Computing Center for providing computational resources essential to this work.

## Notes

### Competing Interest Statement

The authors have declared no competing interest.

